# Mitogenomic analyses provide further evidence for multiple miniaturization events during the evolution of Seychelles caecilian amphibians

**DOI:** 10.1101/2021.03.24.436854

**Authors:** Annie J. Gwilt, Jeffrey W. Streicher, Simon T. Maddock

## Abstract

Whole mitochondrial genomes have been helpful in estimating phylogenetic relationships in many organismal groups, including caecilian amphibians. Despite the increasing ease of obtaining mitochondrial genome sequences from high-throughput sequencing, several species of caecilian lack this important molecular resource. As part of a targeted-sequence capture project of nuclear ultraconserved elements for a small but substantially diverse radiation of caecilian amphibians found on the granitic Seychelles, we examined off-target sequences to determine if we captured enough mitochondrial fragments to reconstruct mitogenomes. We reconstructed (near-)complete mitogenomes for six of the eight species of Seychelles caecilians and completed 14 independent phylogenetic analyses (Bayesian inference and maximum likelihood) on different mitochondrial datasets assembled using different alignment techniques. As with other studies, we were unable to fully resolve internal phylogenetic relationships of the group. However, we found strong support in most analyses that a recently described miniaturized species, *Hypogeophis pti*, and another similarly-sized miniaturized species, *H. brevis* are not sister taxa. Our study suggests that miniature species of caecilians likely evolved at least twice on the Seychelles and highlights the need to revise genus-level taxonomy of Seychelles caecilians while providing further evidence that off-target sequences often contain enough mitochondrial fragments to reconstruct mitogenomes.

## Introduction

Mitochondrial DNA (mtDNA) is a powerful tool used extensively in phylogenetic studies, partly because it is relatively affordable to sequence and because it is found in high copy number in the cell. Although it is still common to utilize single mitochondrial genes, an increasing number of studies are using complete mitochondrial genomes (“mitogenomes” henceforth) to address evolutionary questions. Mitogenomes have been used disproportionately in some organismal groups, for example in the least studied terrestrial vertebrate Order—the Gymnophiona or caecilian amphibians—to estimate robust family level phylogenies (e.g. San Mauro *et al.*, 2014). Whilst mitochondria are circular and are generally considered to lack recombination, some studies of amphibians have found conflicting phylogenetic signals between data (Weisrock *et al.*, 2005; Zhang *et al.*, 2013) and thus utilizing full mitogenomes might provide better estimates of phylogeny than single mitochondrial genes.

Mitogenomic studies are often complemented by studies targeting thousands of regions of nuclear DNA, for example ultraconserved elements (UCEs; Faircloth et al. 2012), due to the vast quantity of data produced and because they are generally unlinked. Targeted nuclear high-throughput sequencing methods can be used to address a multitude of different evolutionary questions (see Zhang, Williams and Lucky, 2019). Due to both imprecise capture and imperfect washing protocols, non-targeted sections of DNA are often sequenced as a by-product. These non-targeted regions of DNA are known as off-target sequences, many of which are short mitochondrial reads due to the high abundance of mtDNA that can be found in each cell (Samuels *et al.*, 2013). Some studies have demonstrated that by using these off-target sequences it is possible to recover not just mitochondrial genes but to reconstruct complete mitogenomes of sequenced organisms (do Amaral *et al.*, 2015; Wang *et al.*, 2017; Zarza *et al.*, 2018; Allio *et al.*, 2020).

The Seychelles is a mid-oceanic archipelago of continental origin, situated in the Indian Ocean. The islands are home to two ancient endemic radiations of amphibians (indotyphlid caecilians and sooglossid frogs) (Nussbaum, 1984), one transoceanic arrived hyperoliid treefrog (Maddock *et al.*, 2014), and one alien introduced frog (Williams *et al.*, 2020). The eight indotyphlid caecilians (Maddock *et al.*, 2018, 2017; Nussbaum, 1984) are distributed on up to 11 of the granitic islands (Maddock *et al.*, 2020): *Praslinia* Boulenger, 1909 (*P. cooperi* Boulenger, 1909)*, Hypogeophis* Peters, 1880 (*H. brevis* Boulenger, 1911; *H. montanus* Maddock, Wilkinson & Gower, 2018; *H. pti* Maddock, Wilkinson, Nussbaum & Gower, 2017; *H. rostratus* (Cuvier, 1829)), and *Grandisonia* Taylor, 1968 (*G. alternans* (Stejneger, 1893); *G. larvata* (Ahl, 1934); *G. sechellensis* (Boulenger, 1911)).

Although seemingly few in number (yet accounting for ca. 4% of the known global caecilian diversity), phylogenetic relationships among the Seychelles caecilians have remained largely unresolved despite repeated efforts to estimate phylogenetic relationships. Almost all studies have recovered *P. cooperi* as the sister taxon to other members of the radiation, and that *G. larvata* and *G. sechellensis* form a sister relationship to each other (Loader *et al.*, 2007; Maddock *et al.*, 2016; San Mauro *et al.*, 2014). Based on the aforementioned molecular studies, the monophyly of both *Grandisonia* and *Hypogeophis* seem unlikely (though support values for any inter-generic relationships are often low).

In this study we extract mitochondrial reads from libraries of UCE data and use the resultant sequences to estimate evolutionary relationships of the Seychelles caecilians. We incorporate newly generated and previously published data, provide data on a heretofore unsampled taxon (*H. pti*) and add to the growing evidence in support of mitochondrial by-products being present in datasets generated from targeted sequence capture and high-throughput sequencing.

## Materials and Methods

Tissue samples (muscle and liver) from nine Seychelles caecilians, representing all of the eight recognized species, were collected in two time periods: 1981–1991 for accession numbers prefaced by RAN (housed at University of Michigan Museum of Zoology); and 2013–2015 for accession numbers prefaced by SM (tissue samples collected by Simon Maddock) (Table 1). Associated specimens for RAN are deposited at the University of Michigan Museum of Zoology, Michigan, USA, and for SM at the Natural History Museum, London, UK. Whole genomic DNA was extracted from tissue samples using a Qiagen DNeasy Blood and Tissue Kit, following manufacturers guidelines. Libraries were prepared for 48 samples, of which nine were Seychelles caecilians, to capture UCEs (Faircloth *et al.*, 2012) following the protocol of Streicher *et al.* (2020) with some minor modifications: fragment sizes of 230–530 bp were selected using a Blue Pippin and paired sequencing was carried out by Novogene China on a single PE150 Illumina HiSeq lane.

**Table 1.**
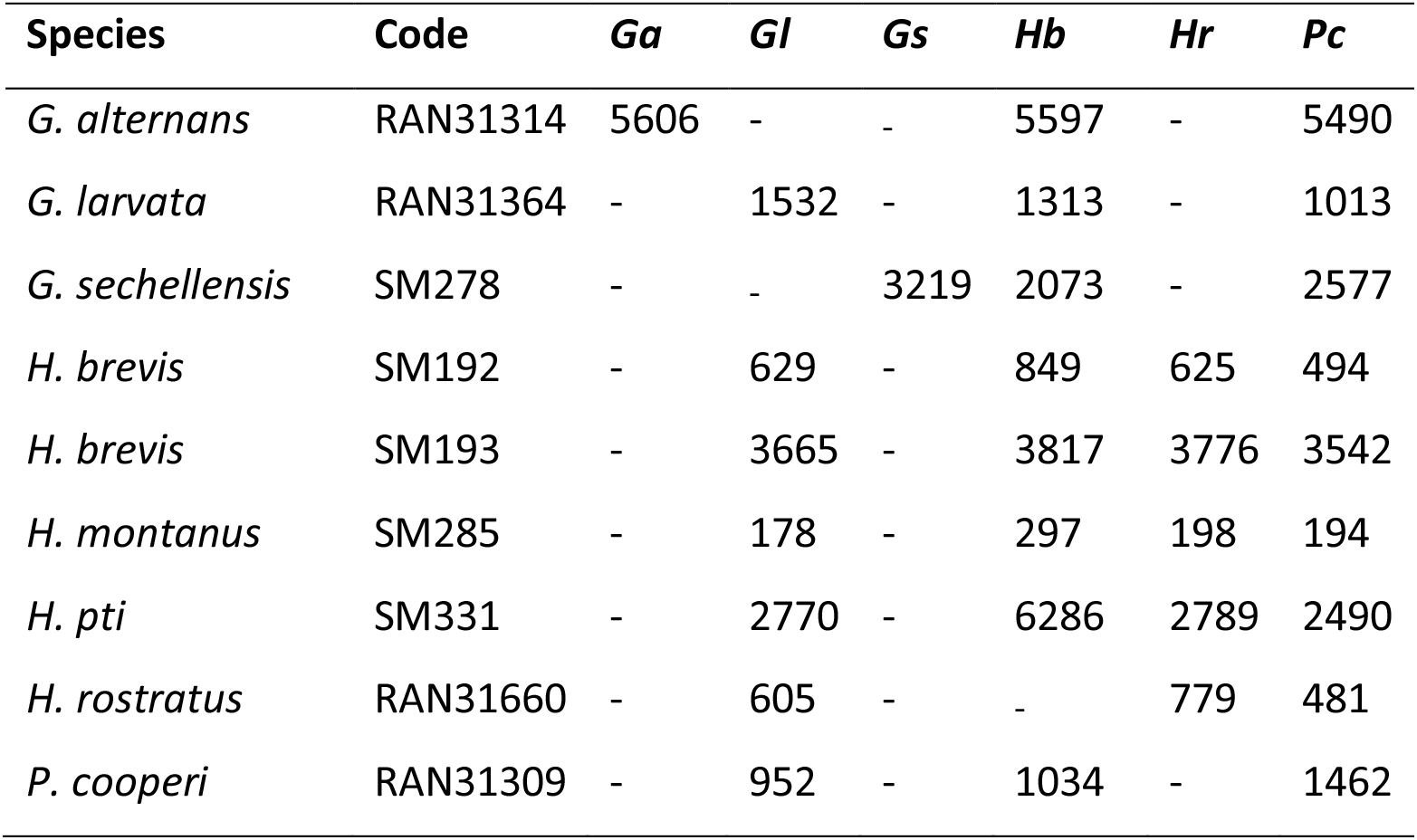
Summary table for number of sequences recovered from each sample when mapped to published mitogenomes using MITObim (see supplementary material for full information about each reconstruction and level of coverage). Only highest numbers of reads displayed for a single reconstruction are displayed. Genbank accession numbers for which samples were mapped were: *Ga* (*Grandisonia alternans*) KU753811, *Gl* (*G. larvata*) KU753813, *Gs* (*G. sechellensis*) KF540152, *Hb* (*Hypogeophis brevis*) KU753817, *Hr* (*H. rostratus*) KF540154, *Pc* (*Praslinia cooperi*) KF540162. Dashes indicate that samples were not mapped to the published mitogenome.

| Species | Code | *Ga* | *Gl* | *Gs* | *Hb* | *Hr* | *Pc* |
| --- | --- | --- | --- | --- | --- | --- | --- |
| *G. alternans* | RAN31314 | 5606 | - | - | 5597 | - | 5490 |
| *G. larvata* | RAN31364 | - | 1532 | - | 1313 | - | 1013 |
| *G. sechellensis* | SM278 | - | - | 3219 | 2073 | - | 2577 |
| *H. brevis* | SM192 | - | 629 | - | 849 | 625 | 494 |
| *H. brevis* | SM193 | - | 3665 | - | 3817 | 3776 | 3542 |
| *H. montanus* | SM285 | - | 178 | - | 297 | 198 | 194 |
| *H. pti* | SM331 | - | 2770 | - | 6286 | 2789 | 2490 |
| *H. rostratus* | RAN31660 | - | 605 | - | - | 779 | 481 |
| *P. cooperi* | RAN31309 | - | 952 | - | 1034 | - | 1462 |

Raw sequences were initially quality checked, trimmed and cleaned using illumiprocessor (Faircloth 2013). MITObim V.1.9.1 (Hahn *et al.*, 2013) was used to reconstruct mitogenomes from the paired reads using the --quick option. Each interleaved file was mapped against a previously published mitogenome (obtained from GenBank) reference sequences from each Seychelles caecilian genus (see Supplementary Material for details). For each sample and mapped genus, mitogenome reconstructions were completed using k-mer sizes of 20, 31 and 45. In addition, we attempted to reconstruct the mitogenome from each sample using the *de novo* assembly method with k-mer sizes of 45.

Contig files produced by MITObim were imported into Geneious Prime 2020.1.2 for further investigation and consensus sequence construction. MITObim analyses for each sample were combined together by using the ‘Map to Reference’ function using default settings to the assumed closest relative. A consensus sequence was created, taking into account all reconstructions and imported into the MITOS Web Server (Bernt *et al.*, 2013) for annotation. Annotated mitogenomes were imported into Geneious and aligned to other Seychelles caecilian mitogenomes (obtained from GenBank) using the Geneious Alignment method. Manual editing of mitogenomes then took place considering the MITOS results and annotated mitogenomes from GenBank; this primarily involved ensuring that the end of a protein coding gene contained a stop codon.

To remove poorly aligned positions and/or divergent regions of DNA, the GBlocks Server v 0.91b (Castresana, 2000) was used, with settings set to allow gaps and smaller blocks. All subsequent analyses, where gaps were previously present (tRNA and rRNA data), were analyzed for the pre- and post-GBlocks datasets. PartitionFinder V.2.1 (Lanfear *et al.*, 2017) was used to determine the optimum partitioning strategies (for maximum likelihood (ML and Bayesian inference (BI)) and the best models of molecular evolution (for BI) for analyzing the data for seven datasets: (1) protein-coding genes, (2) tRNA pre-GBlocks, (3) tRNA post-Gblocks, (4) rRNA pre-GBlocks, (5) rRNA post-GBlocks, (6) all data pre-GBlocks, and (7) all data post-GBlocks. We removed the regulatory, non-coding L-strand replication and control regions from the alignment prior to phylogenetic analyses following San Mauro *et al.* (2014) and Maddock *et al.* (2016).

All phylogenetic analyses were run through the CIPRES Science Gateway V3.3 (Miller *et al.*, 2010). For each of the datasets, ML trees were reconstructed using RAxML V.8.2.12 (Stamatakis, 2014) and run for 500 bootstraps using the GTRGAMMA model. Bayesian Inference trees were estimated using MrBayes V.3.2.7a (Ronquist *et al.*, 2012) and run for 10^7^ generations for one hot and three cold chains. BI trees were sampled every 1,000 generations, with a 10% burnin. Tracer V1.7.1 (Rambaut *et al.*, 2018) was used to evaluate convergence by assessing that effective sample size (ESS) values were >200 and that the potential scale reduction factor (PSRF) was close to 1.0. *Praslinia cooperi* was used as the outgroup for all analyses following the general consensus that it is the sister species to all other Seychelles caecilians (Loader *et al.*, 2007; Maddock *et al.*, 2016; San Mauro *et al.*, 2014).

## Results and Discussion

The UCE data allowed for the reconstruction of (near-)complete mitogenomes for seven of the nine samples sequenced in this study; the two remaining samples (SM285 *Hypogeophis montanus* and RAN31660 *H. rostratus*) were not included in phylogenetic analyses due to the small amount of mitochondrial data recovered from them (Table 1). The successful reconstruction of a mitogenome from *H. pti* represents the first for the species and out of the eight currently recognized Seychelles caecilian species, only *H. montanus* lacks a sequenced/reconstructed mitogenome.

Whilst mitogenome recovery was not successful for all samples, our results provide growing support that UCE datasets often contain large quantities of off-target mitochondrial reads (do Amaral *et al.*, 2015; Wang *et al.*, 2017; Zarza *et al.*, 2018; Allio *et al.*, 2020). The mean sequencing depth for all mitogenomes, under each mitogenome recovery step, averaged 14.9 (minimum 1.4 and maximum 56.1).

A k-mer value of 20 always outperformed other k-mer reconstruction values except when *Praslinia cooperi* was mapped to the published *P. cooperi*, which resulted in all reconstructions recovering the same numbers of reads (Supplementary Table 1). *De novo* assemblies, not to a reference mitogenome, resulted in poor mitogenome recovery.

Of the 14 different phylogenetic analyses in this study, five consensus topologies were estimated (Fig. 1). In all analyses, *Grandisonia larvata* and *G. sechellensis* were recovered as sister species with maximal support (bs = 100; PP = 1.0), apart from in the rRNA and tRNA ML analyses, but the relationship was still supported with minimum bootstrap posterior probabilities of 90. The sister relationship between *G. larvata* and *G. sechellensis* has previously been recovered (Loader *et al.*, 2007; Maddock *et al.*, 2016; San Mauro *et al.*, 2014). Ten of the 14 trees maximally, or strongly, support *H. pti* being the sister species to the *G. larvata + G. sechellensis* clade. The four analyses that do not support the (*H. pti* (*G. larvata + G. sechellensis*)) relationship (Fig. 1d) are all estimated using the less phylogenetically informative tRNA data, which is also the dataset with the shortest alignment, for which most internal relationships within the tree are poorly resolved.

**Fig.1.**
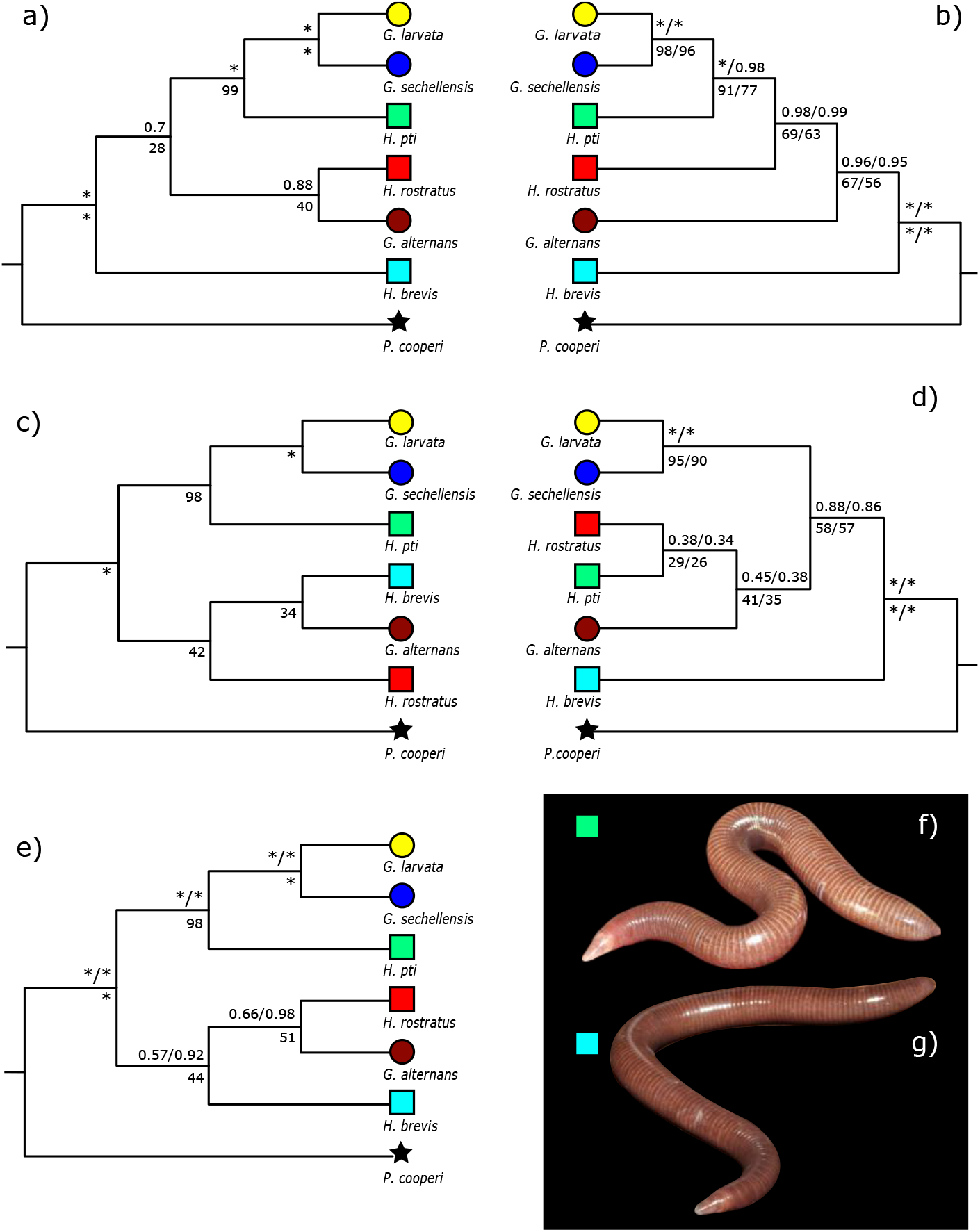
Evolutionary relationships generated from 14 phylogenetic analyses representing five differing topologies. In all phylogenetic trees, numbers above branches are Bayesian posterior probabilities and numbers below branches are for ML support estimates. Full support is indicated with a “*”. Phylogenetic topologies refer to: **(a)** all data pre-GBlocks (BI+ML), **(b)** rRNA pre- (BI+ML) and post-GBlocks (BI+ML), **(c)** all data post-Gblocks (ML), **(d)** tRNA pre- (BI+ML) and post-Gblocks, **(e)** all data post-Gblocks (BI) and protein coding DNA (BI+ML). Images in life of **(f)** *H. pti* and (g) *H. brevis.* Symbols indicate genus: star = *Praslinia*, square = *Hypogeophis* and circle = *Grandisonia.* Colours refer to species: Black = *P. cooperi*, red = *H. rostratus*, turquoise = *H. brevis*, green = *H. pti*, brown = *G. alternans*, blue = *G. sechellensis* and yellow = *G. larvata*.

Relationships among the remaining three internal taxa (*H. brevis, H. rostratus* and *G. alternans*) largely remain uncertain, as found in previous studies (Loader *et al.*, 2007; Maddock *et al.*, 2016; San Mauro *et al.*, 2014). The missing taxon for which we were unable to reconstruct a mitogenome (*H. montanus)* might help resolve these internal relationships by shortening branch lengths. Morphological and molecular support exists for *H. montanus* being the sister taxon to *H. brevis* (Maddock *et al.*, 2018). Similar to the findings of Maddock *et al.* (2016) it is noteworthy that *H. brevis* and *H. rostratus* never form a sister relationship in any of our topologies.

That *Grandisonia* and *Hypogeophis* seem to lack generic level monophyly with respect to one another suggests that the genera may need a taxonomic review. Increasing the size of phylogenetic datasets by adding *H. montanus* to the mitogenome phylogeny, by accounting for intraspecific variation (e.g., such as that found within *H. rostratus* (Maddock *et al.*, 2020), or by incorporation of a multi-gene nuclear dataset, may help better resolve the relationship of the Seychelles caecilians and provide support for a generic level revision.

Interestingly, that *H. brevis* and *H. pti* are almost certainly not sister taxa, suggests a possible convergence of body form: miniaturized, heavy set and having elongated, pointed snouts. Little is known about the ecologies of the species, but this possible convergence may indicate adaptation to similar niches on their respective islands: *H. brevis* inhabits Mahé and *H. pti* inhabits Praslin. Based on limited data, Maddock, Wilkinson and Gower (2018), also hypothesized about this possible convergence of body form between *H. brevis + H. montanus* vs. *H. pti*. The convergence of the elongated and pointed snout shape is particularly intriguing, considering that this feature does not appear in any other known caecilian.

Our study builds on previous evidence (do Amaral *et al.*, 2015; Wang *et al.*, 2017; Zarza *et al.*, 2018; Allio *et al.*, 2020) that a large number of off-target reads from targeted sequence capture are mitochondrial. These data often have phylogenetic or other (e.g. metabolic study) value that could and should be exploited. The support from mitogenomes for the convergence of smaller-than-ancestral body sizes in at least two lineages of Seychelles caecilians will be important to compare to the complimentary nuclear UCE analyses that will be possible with the analyzed dataset.

## Acknowledgements

We thank Gregory Schneider and Ronald Nussbaum for providing access to tissue samples from UMMZ. The Seychelles Bureau of Standards for providing required field collection permits; and the Seychelles National Park Authority for supporting STM with a gainful occupancy permit. The Mohammed Bin Zayed Conservation Fund financially supported fieldwork. We thank all project partners for support throughout the project and the Seychelles Ministry of Environment for issuing permits for research and export. In addition, we thank David Gower, Jim Labisko and Mark Wilkinson for assistance with field collection of samples.

